# Network reorganization distinguishes vulnerability and resilience to observational fear

**DOI:** 10.64898/2026.03.12.711378

**Authors:** Nicola Murgia, Botond Molnár, Janine Reinert, Kanat Chanthongdee, Antonio Lacorte, Li Xu, László Mihály, Mária Ercsey-Ravasz, Christoph Körber, Estelle Barbier

## Abstract

Individuals vary widely in their responses to stress and threat, with some developing persistent fear after adverse experiences while others remain resilient. Such variability also extends to social contexts, where individuals can acquire information about danger by observing others in distress through observational fear learning. The neural mechanisms underlying individual differences in responses to socially conveyed threat remain poorly understood. Here, we examined how variability in OFL relates to large-scale brain network organization. Rats observed conspecifics receiving tone-shock pairings and were later tested for fear responses to the conditioned stimulus. Behavioral analysis revealed two phenotypes: observational-susceptible rats displaying robust freezing and observational-resilient rats showing freezing levels comparable to controls. Despite these differences, both groups exhibited elevated corticosterone responses, indicating that socially conveyed threat was detected across animals. Brain-wide c-Fos mapping across 84 regions combined with graph-theoretical analysis revealed distinct network architectures associated with each phenotype. These findings suggest that susceptibility and resilience to socially acquired fear emerge from differences in distributed brain network organization.

## INTRODUCTION

Individuals vary widely in how they respond to adverse experiences. While some develop persistent fear and anxiety after stress exposure, others remain resilient despite encountering similar challenges (*1*). Understanding the neural mechanisms of individual differences is crucial for elucidating the biological bases of both vulnerability and resilience in anxiety- and trauma-related disorders (*2*). However, the neural mechanisms that bias individuals toward maladaptive versus resilient response remain poorly understood.

In social species, fear information is often acquired indirectly by observing others in distress (*3*). This process, termed observational or vicarious fear learning, allows individuals to anticipate danger without direct exposure, enhancing survival. However, repeated, or intense exposure can promote persistent fear and anxiety, and has been linked to elevated risk for stress-related psychopathology(*3–6*). For example, evidence from war veterans indicates that socially transmitted fear can exert profound psychological effects, sometimes rivalling or exceeding those of direct trauma (*4*). Similarly, professionals with chronic occupational exposure to others’ trauma, such as firefighters and trauma nurses, exhibit higher rates of anxiety disorders and PTSD (*6*). Importantly, these effects are not restricted to high-risk populations; comparable associations have been reported in the general population following repeated exposure to traumatic events via social media (*5*). Together, these observations highlight the broad relevance of socially acquired fear and motivate its investigation as a conserved learning process that contributes to vulnerability to stress-related psychopathology.

Socially acquired fear can be studied using observational fear learning (OFL), a process that is conserved across species, including humans, non-human primates, and rodents (*3, 7, 8*). In both rodent and humans, observational fear engages neural circuits that partially overlap with those involved during direct fear learning, including the amygdala, anterior cingulate cortex, and insula (*3, 9–11*). These studies have substantially deepened our understanding of the mechanisms through which fear memories are encoded and subsequently consolidated. However, their group-level design fails to capture individual variability, providing only limited insight into how neural processes are organized to produce distinct behavioral outcomes. In contrast, examining individual differences can uncover the processes that promote vulnerability to anxiety and trauma-related disorders, rather than only characterizing broad aspects of fear memory processing.

Cognitive functions, including emotional fear responses, are widely thought to depend on interactions among distributed brain regions, with specific circuits embedded within broader network organization (*12*). We hypothesized that variability in fear responses may be associated with differences in coordinated activity across large-scale brain networks supporting emotional processes, which in turn may influence the function of established fear circuits.

Activity mapping across larger brain areas combined with network analysis provides a powerful framework for uncovering how functional architecture reorganizes in response to socially acquired fear, and for determining whether such reorganization differs between susceptible and resilient individuals. Here, we combine behavioral phenotyping with whole-brain cFos mapping and graph-theoretical analysis in a rat model of OFL to examine how large-scale network organization relates to individual variability in responses to socially acquired fear. Observer rats segregate into two behavioral phenotypes that differ markedly in defensive responses despite comparable endocrine activation. We show that these phenotypes are associated with distinct brain-wide network architectures, including differences in mesoscale clustering, hub organization, and communication efficiency. These findings provide a systems-level framework for understanding how socially acquired fear is represented across distributed brain networks.

## MATERIAL AND METHOD

### Animals

Adult male Wistar rats (250–300g; Charles River, Germany) were pair-housed with a same-sex conspecific in a humidity- and temperature-controlled environment with food and water access ad libitum under a reverse light cycle. Rats were acclimated to the facility and handled by experimenters prior to behavioral experiments. To familiarize rats with their cage-mates, rats were housed in pairs with a same-sex conspecific for at least 4 weeks. All experiments took place during the dark phase. Procedures were approved by the Local Animal Ethics Committee at Linköping university (Dnr 16869-2022 ID 4498) and were in accordance with the EU Directive 2010/63/EU on the protection of animals used for scientific purposes as implemented in Swedish national regulations.

### Apparatus

All experiments took place in operant chambers (25 cm x 32 cm x 26 cm, Med Associates Inc., Fairfax, VT, USA) equipped with two retractable levers, rod flooring connected to a shock generator, and a speaker inside an individual sound-attenuated cubicle. The operant chambers were placed in sound-attenuating cubicles. A custom-built transparent perforated divider was installed inside each chamber, and a black floor was added exclusively to one side of the chamber during observational fear conditioning to ensure that observer rats did not receive the footshock. Rats were tested for fear memory in a smaller operant chamber (25 cm x 16 cm x 26 cm). All chambers were controlled by MEDPC-IV software (Med Associates Inc.)

### Behavioral procedures

#### Observational fear conditioning and fear testing

In each dyad, one rat was designated as the demonstrator and the other as the observer. The observational fear conditioning protocol lasted five days. On Days 1 and 2, observer rats were habituated for 20 minutes/day to the floor and the chamber to reduce distraction from a novel context. Our pilot experiment and previous findings suggested that conditioned observational fear in rodents is promoted by prior shock-experience (*7*). Therefore, on Day 3, rats in the observer group were exposed to four unpredictable trials of 0.8 mA electric footshocks (3 minutes + 4 x 2 seconds shock (10-80 seconds inter-trial interval) + 3 minutes) in a different context (no illumination, different room). On Day 4, observers and demonstrators rats underwent observational fear conditioning in a two-compartment chamber with a transparent perforated divider, allowing visual contact and interactions by sniffing. After 5 minutes of initial habituation, demonstrators were exposed to six trials of conditioned stimuli (CS; 30 seconds, 2.9 kHz pure tone, 80 dB; 3 minutes inter-trial interval) that co-terminated with 2 seconds 0.8 mA electric footshocks. During conditioning, observer rats were placed on the black floor on the other side of a divider and were able to witness demonstrator rats receiving electric footshocks at the end of the tones. Unless specified otherwise, on Day 5, all rats individually underwent fear memory testing. After 5 minutes of habituation, two trials of 30s CS with a 3 minutes inter-trial interval was presented in the absence of electric shocks.

Fear expression was quantified as the percentage of time spent freezing during CS presentations and was scored by two trained experimenters or automatically using EthoVision XT17 (cFos experiment). In all cases, the experimenters performing the scoring were blind to the animals’ group identity. ‘Freezing’ is defined as an absence of any movements except respiration that last longer than one second (Blanchard and Blanchard, 1969). Rats were classified based on their freezing responses during tone presentation. Animals exhibiting ≥10 % freezing were categorized as observational-susceptible (OBS-S), whereas animals showing no freezing responses were categorized as observational resilient (OBS-R). This threshold was chosen because baseline freezing in control animals was minimal (mean ≈ 0.5 %), ensuring that ≥10 % freezing represents a robust fear response well above spontaneous immobility levels.

#### Ethological analysis of behaviors during observational fear testing

To quantify a broad range of behaviors during the expression of vicarious fear, video recordings obtained during the 30-s conditioned stimulus (CS) presentations in the fear test were analyzed for time spent engaging in observable behaviors using the manual scoring function in EthoVision software (Version 17, Noldus, Amsterdam, The Netherlands), as previously described (Chanthongdee et al., 2024). Time spent performing each behavior was calculated as a percentage of the scoring period.

#### Plasma corticosterone analysis

Blood samples were collected from tail veins into heparin-coated tubes and centrifuged for 10 minutes at 5,000 x g to separate plasma (*13*). Plasma was transferred into new tubes and stored at –80 °C for further analysis. Corticosterone was extracted by adding five parts of ethyl acetate (Thermo Fisher Scientific Inc. Waltham, MA, USA) to each plasma sample. The organic solvent layer was first transferred to a water-prefilled tube and then to the second tube. This procedure was repeated two times before samples were dried in a vacuum concentrator. Samples were re-dissolved in assay buffer from the DetectX Corticosterone Enzyme Immunoassay Kit (Arbor Assays, Ann Arbor, MI, USA). All samples were then processed following the manufacturer’s instructions.

#### Brain collection and cFos immunohistochemistry

Ninety minutes after observational fear testing, observers, together with control observers (Obs-Ctrl), were anaesthetized with isoflurane (3–4 %) and transcardially perfused with phosphate-buffered saline (PBS, pH 7.4) followed by 4 % paraformaldehyde (PFA). Brains were collected and stored in PBS before being flash-frozen using isopentane. Coronal sections (30 µm thick) were collected with a Epredia CryoStar NX50 cryostat and stored free-floating in cryoprotectant (1X PBS containing 30 % ethylene glycol and 20 % glycerol) at –20 °C. Free-floating brain sections were processed for c-Fos immunofluorescence. Briefly, on Day 1, sections were removed from the antifreeze solution and rinsed in 1× PBS (3 × 10 min, room temperature). Sections were then incubated for 60 min in blocking solution containing 4 % BSA in PBS-T (0.2 % Triton X-100 in 0.1 M PBS), followed by overnight incubation at 4 °C in a humid chamber with gentle agitation in rabbit anti-c-Fos primary antibody (Invitrogen MA5-15055; 1:1000) diluted in 4 % BSA in PBS-T. On Day 2, sections were washed in 1 x PBS (3 × 10 min) and incubated for 1.5 h at room temperature with donkey anti-rabbit Alexa Fluor 488 secondary antibody (Thermo Fisher A-21206; 1:500) diluted in PBS-T, after which all steps were performed in the dark. Slices were washed again in 1 x PBS (3 × 10 min) and incubated for 30 min at room temperature with Hoechst 33258 (Thermo Fisher Catalog number H3569; 1 µg/ml in PBS). After a final series of PBS washes (3 × 10 min), sections were mounted on Superfrost™ Plus slides and coverslipped. Fluorescent imaging was performed in three rounds to cover the targeted brain regions. In the first round, sections at +3.0 and +2.7 mm from bregma were imaged to capture the infralimbic (IL), dorsal peduncular, and prelimbic (PrL) cortices, and sections at +2.2 and +1.8 mm were used to image the nucleus accumbens (NAc) and lateral septum (LS). The second round included sections containing the bed nucleus of the stria terminalis (BNST), as well as sections at −1.2, −1.4, and −1.5 mm for imaging the paraventricular thalamic nuclei (PVA and PVT). In the third round, sections at −2.4 and −2.7 mm were imaged to analyze the hypothalamic, amygdala, lateral habenula, and zona incerta regions.

#### Image acquisition and analysis

Images of complete stained brain sections were assembled from tile scans acquired using an epifluorescence microscope (Zeiss Axio Observer.Z1/7, Zeiss, Germany) equipped with a 10x objective (0.3 NA, M27 EC-Plan-Neofluar, Zeiss, Germany, image resolution: 0.586 µm x 0.586 µm x 1.0 µm). Images were registered to the Waxholm space atlas (*14*) using the QUINT pipeline (*15*). In brief, the Hoechst channel of each image was used for alignment to the reference atlas while both the Hoechst and cFos channel were subsequently used to determine the number of cFos-expressing cells per reference brain region. Pre-processing of the images was performed using FIJI (*16*) (ImageJ 1.54f, Java 1.8.0_322). For registration, images were filtered (Top Hat at radius = 20 pixel followed by Gaussian Blur filters at sigma = 1) and down-sampled to a width of 600 pixels. For quantification, images were filtered similarly (Hoechst channel: Top Hat at 20 pixel radius; cFos channel: Top Hat at 15 pixel radius; both channels: Gaussian Blur at sigma = 1) and used at full resolution for object classification using separately trained classifiers (Ilastik; (*17*)). Co-localization of Hoechst and cFos objects was determined based on the Euclidian distance of the object centres using custom written Python routines (Python 3.9). Only objects with a minimal size of 60 pixels (approx. 16 µm diameter) were included. Objects whose centers were located at less than 20 pixels (approx. 11 µm) apart were considered as colocalized. The amount of cFos expression per brain region was determined as the ratio between objects showing colocalization of Hoechst and cFos and the number of Hoechst-positive objects. Moreover, cFos expression was analyzed in a total of five brain regions that were not included in the Waxholm reference atlas: basolateral amygdala (BLA), central amygdala (CeA), medial amygdala (MeA), perifornical lateral hypothalamus (PeFLH), and ventromedial nucleus of the hypothalamus (VMN). These regions were manually delineated based on the Hoechst signal and quantified using the QUINT pipeline described above.

#### Network Analysis

To analyze the characteristics and quantify the changes of the experimental data, first we have constructed a global brain network for each experimental group. Edge weights were defined as Pearson correlation coefficients values calculated between brain regions across the cohorts of each experimental group. This yielded a weighted, undirected, and fully connected graph. To preserve the full topological structure of the network, the analysis was conducted without thresholding or the elimination of weak links. Given that the weights represent correlation coefficient values, they range between [−1, 1]. Let 𝑤_𝑖𝑗_ denote the weight of the link between nodes 𝑖 and 𝑗. In the current study, we focus our study on the clustering of the Pearson correlation network, the backbone, communication efficiency and various centrality measures, including betweenness, closeness and weighted degree centrality, as well as the Hamiltonian path.

**Voronoi clustering** was developed as a new approach for identifying optimal groups within networks, inspired by the Voronoi diagrams in the Euclidian space and later generalized for weighted, directed networks (*18*). In the classical geometric sense, a Voronoi diagram partitions a space into regions based on distance to a specific set of so-called generator points. In the context of a brain network, this concept is adapted by using functional distance (defined by the shortest path) instead of physical distance. The distances between nodes were defined as 𝑙_𝑖𝑗_ = − ln|𝑤_𝑖𝑗_|. The clustering process follows two main steps:

**Generator point selection:** A set of nodes is selected as seeds or cluster centers by calculating the local relative density of the nodes considering the strength of a node and the number of outgoing and incoming links of the first order neighborhood of a node. It is crucial to accurately define these generator points, because they represent the core regions around which functional communities are organized.

**Cell Assignment:** Each remaining node in the network is assigned to the cluster of the generator point to which it has the shortest weighted path. This effectively partitions the network into influence zones where all regions within a cluster are functionally closer to their own generator point rather than to any other seed in the network.

To find the optimal clustering, we vary the radius (a resolution parameter) to identify the configuration which yields the highest modularity. This approach is particularly useful in neuroscience as it does not require an a priori assumption about the number of clusters and accounts for the specific bandwidth of information flow between regions.

**Centrality measures** identify the hub nodes in a network. In neuroscience, hubs (regions with high centrality values) are the most metabolically expensive and the most vulnerable to disease or experimental manipulation. Comparing these across groups shows if the experimental condition shifted the importance from one brain area to another. In the current study we focused on three centrality measures: weighted degree (strength); closeness and betweenness centralities. It is imperative that these measures account for the weights of the links.

**Weighted degree (strength).** In a fully connected graph, the simple degree (the number of connections) is identical for each node (𝑘_𝑖_ = 𝑁 − 1). Therefore, we use the strength (𝑠_𝑖_), defined as the sum of the absolute value of the link weights connected to node 𝑖:

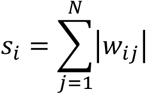

High strength indicates a brain region directly connected to many others with strong connections (whether positive or negative). Intuitively, a node with high strength represents a “social” hub with numerous strong relationships.

**Closeness centrality** measures the proximity of a node to all other nodes in the network. It is defined as the inverse of the average distance from node 𝑖 to all other nodes. A node with high closeness centrality marks a brain region that is functionally closer to the rest of the network, allowing for a faster information exchange.

**Betweenness centrality** quantifies how many shortest paths between all possible pair of nodes pass through the node of interest. In this study, the shortest paths are calculated using the absolute values of weights. To accurately map the weights to topology, strong links (high weight) are treated as short distances, while weak links represent long connections. Following the approach used in anatomical brain networks (*19*) the length of a link is defined as 𝑙_𝑖𝑗_ = − ln|𝑤_𝑖𝑗_|. As edge weights can be considered as proxy for the bandwidth of information flow, the probability of information traveling from node 𝑖 to 𝑚 through node 𝑗 would be proportional to 𝑤_𝑖𝑗_ 𝑤_𝑗𝑚_. Using the logarithmic form makes these values additive, ensuring that shortest paths indicate routes where information flows with the highest probability. A node with high betweenness centrality acts as a broker, facilitating communication between different regions/lobes of the brain.

**Communication efficiency** quantifies the capacity of a network for information processing and exchange. We evaluated this through two distinct measures: global and local communication efficiency. Global efficiency (𝐸_𝑔_) describes the ability of a network to transfer information across the entire system. It is proportional to the average of the inverse resistance (here defined as the length of shortest path) between all node pairs:

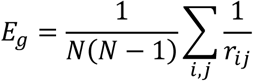

Local efficiency (𝐸_𝑙_), conversely, measures the fault tolerance of the network by assessing the communication efficiency within the neighborhood of each node. It indicates how well information is shared among the neighbors of a specific region if that node and all its links were removed. The local efficiency for the entire network is averaged over all nodes:

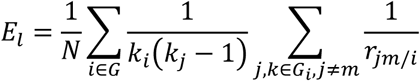

indicates the set of neighbors of node 𝑖; 𝑗 and 𝑘 are neighbors of 𝑖, 𝑟_𝑗𝑘/𝑖_ is the shortest path between 𝑗 and 𝑘 after the removal of node 𝑖 and its links from the graph; 𝑘_𝑖_represents the degree (number of neighbors, links) of node 𝑖.

To further characterize the robustness of these communication pathways, we do not simply calculate these measures but determine their values at different densities of the network, by employing two link removal approaches. We systemically extract the links according to their strength and recalculate both measures for the networks obtained:

**weak link removal:** Links are removed in increasing order of weight (from weakest to strongest correlation). This approach identifies the core or the backbone of the network and tests whether global communication is maintained by the most dominant functional connections.

**strong link removal:** Links are removed in decreasing order of weight (from strongest to weakest). This procedure assesses the vulnerability of the network to the loss of its most significant connections, revealing whether the system relies on the strong ties or possesses redundant, weaker pathways that can sustain communication.

We continue the removal of the links in both cases until the network is completely depleted of links.

**The Backbone** of a network is defined as the core subgraph that connects all nodes in the network, while minimizing the number of edges. To obtain this structure, we employed a process where the network is diluted by removing the links in increasing order of their weights (from the weakest to the strongest correlation). to gradually remove the weakest links one-by-one from the network and stop right before it gets disconnected (falls apart). This resulted in the most robust pathways of communication within the network. Within the context of functional neuroscience, the backbone represents the invariant structure of the network. By removing links until the network is about to disconnect, one can isolate the critical infrastructure that maintains global communication. It ensures that the differences between groups are based on the primary pathways, not random fluctuations in weak correlations.

## RESULTS

### Behavioral phenotyping following observational fear learning

Rats were classified based on freezing responses during tone presentation, with animals exhibiting ≥10% freezing categorized as observational-susceptible (Obs-S) and animals showing no freezing categorized as observational-resilient (Obs-R). Among the 167 observer rats used for behavioral classification, most were categorized as Obs-R (∼82%), whereas a smaller subset displayed robust freezing responses and were classified as Obs-S (∼18%) (Fig. 2A). An additional group of 18 control observer rats (Obs-Ctrl) were included for comparison. Individual freezing values confirmed that Obs-S rats exhibited markedly higher freezing than both Obs-R and Obs-Ctrl rats, whereas Obs-R rats showed freezing levels comparable to controls (Fig. 2A, D).

**Figure 1:**
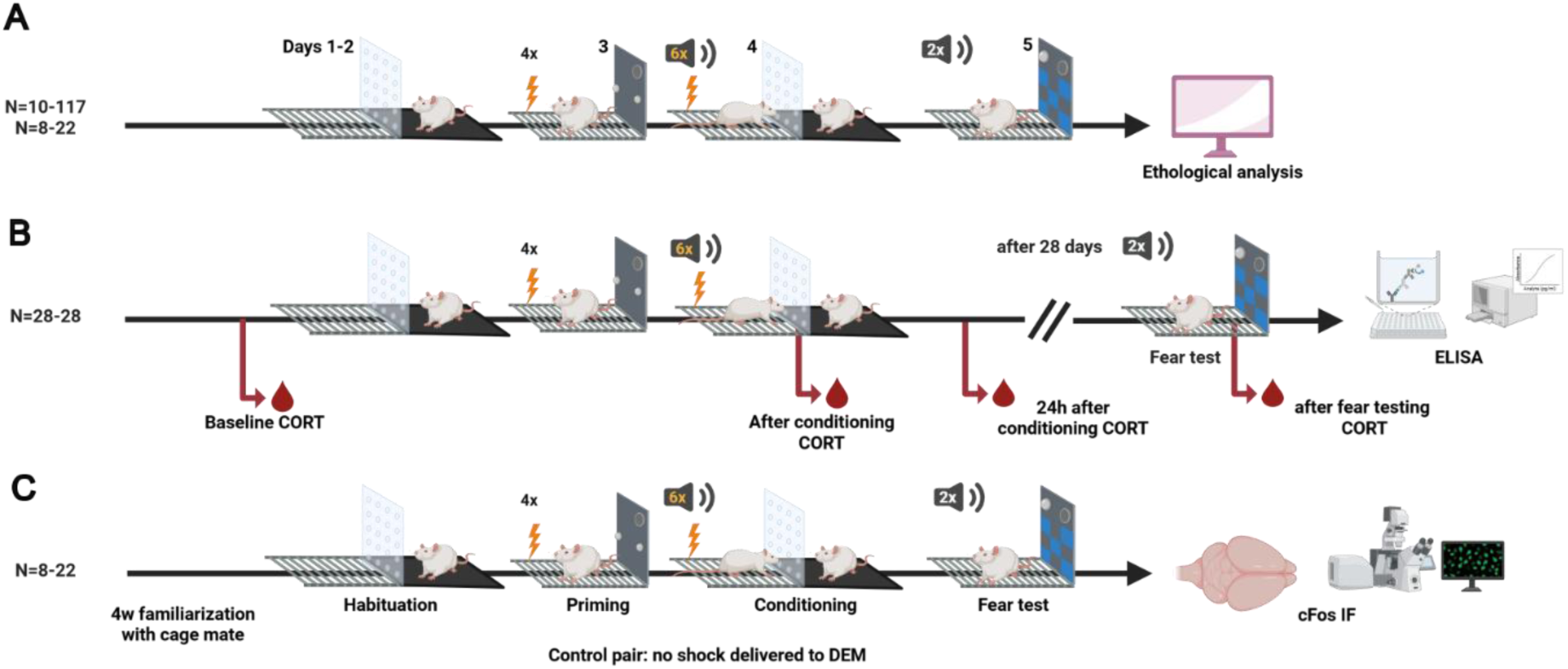
Experimental design and behavioral phenotyping following observational fear learning (OFL). (A) Observational fear learning paradigm. Observer rats were placed in an adjacent compartment separated by a transparent perforated divider and exposed to demonstrators receiving six tone-shock pairings. Observers were later tested for fear expression during tone (CS) presentations. (B) Behavioral phenotyping and endocrine measurements. Observer rats were classified according to freezing responses during CS presentations. Blood samples were collected at multiple time points for corticosterone measurement. A subset of animals underwent a delayed fear test (∼28 days). (C) Brain activity mapping. A separate cohort of observer rats underwent the OFL paradigm and was sacrificed following retrieval for c-Fos immunohistochemistry. c-Fos expression was quantified across 81 brain regions, and correlation matrices were used to construct functional brain networks for graph-theoretical analyses.

**Figure 2:**
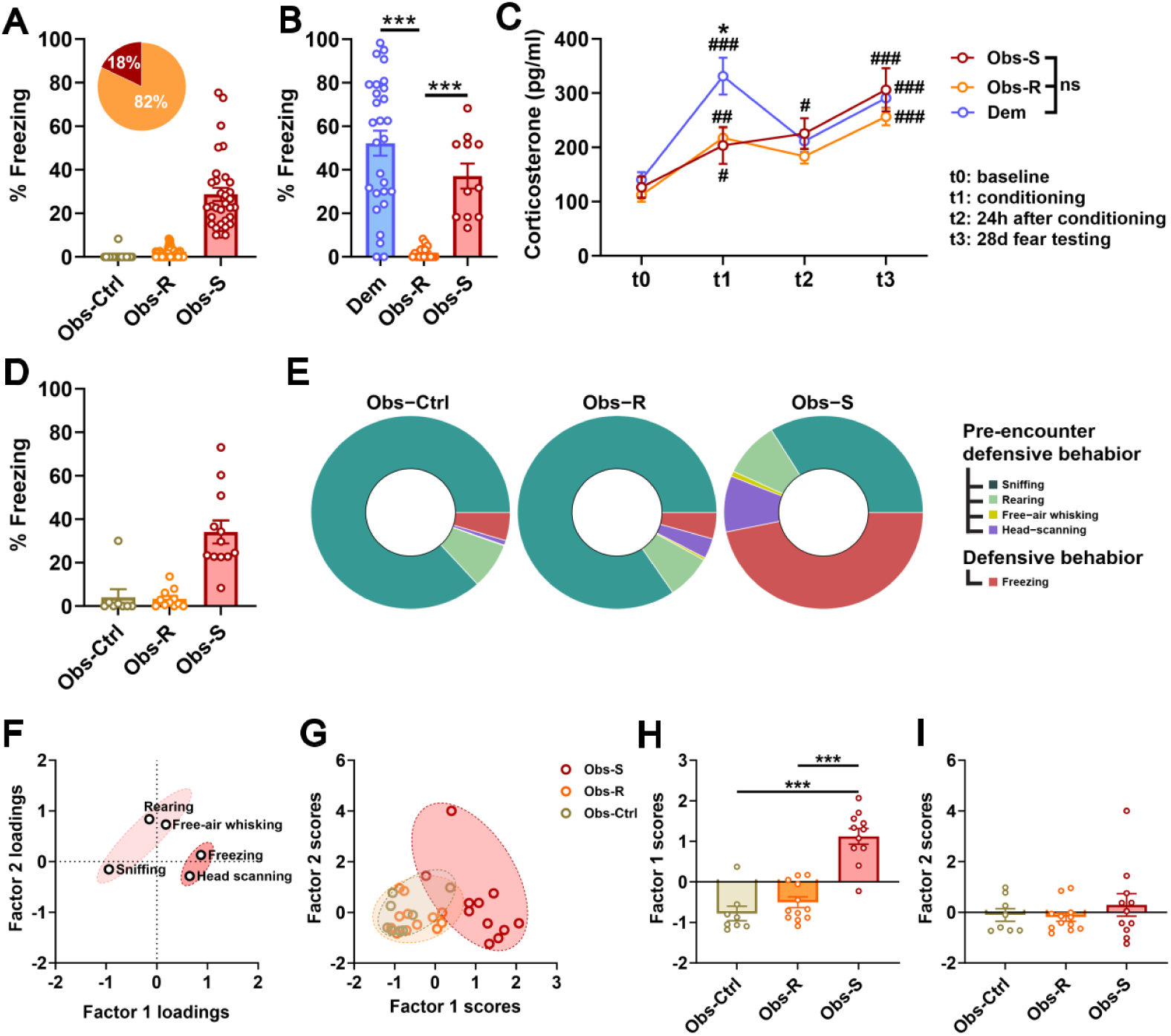
Behavioral phenotyping and ethological characterization following observational fear learning (OFL). (A) Distribution of behavioral phenotypes among observer rats based on freezing responses during conditioned stimulus (CS) presentations. Animals were classified as observational-resilient (Obs-R) or observational-susceptible (Obs-S). (B) Freezing responses during CS presentations across groups. (C) Plasma corticosterone levels measured during OFL acquisition and retrieval. (D) Freezing responses during fear testing. (E) Ethological analysis showing the proportion of time spent in exploratory and defensive behaviors across groups. (F) Factor analysis of ethologically scored behaviors. (G) Clustering of animals based on behavioral factor scores. (H–I) Group comparisons of behavioral factor scores. Data are shown as mean ± SEM with individual animals represented by dots.*p<0.5 compared to demonstrators; #p<0.05 compared to baseline corticosterone.

To determine whether socially transmitted threat information was detected across animals, plasma corticosterone levels were measured during OFL acquisition and retrieval. Both Obs-S and Obs-R rats showed elevated corticosterone responses compared with control observers, indicating that the OFL experience was biologically salient across animals despite differences in defensive behavior (Fig. 2C). Importantly, corticosterone levels did not differ significantly between Obs-S and Obs-R rats, suggesting that both groups registered the socially conveyed threat even though only Obs-S animals displayed overt freezing during CS presentations.

To examine neural network organization associated with OFL-induced behavioral phenotypes, we selected a subset of observer rats for whole-brain cFos mapping (n = 30; 8 Obs-Ctrl, 12 Obs-S, and 10 Obs-R). Before neural analyses, behavioral features in this cohort were further characterized using automated ethological tracking (EthoVision) to obtain a more detailed description of behavior during fear expression beyond freezing alone. Behavioral ethograms showed that Obs-Ctrl and Obs-R rats spent most of the pre-encounter period engaged in exploratory behaviors such as sniffing and rearing, whereas Obs-S rats displayed a substantially higher proportion in defensive behaviors such as freezing behavior and head scanning (Fig. 2E).

Factor analysis of ethologically scored behaviors identified two principal components explaining 69.6% of the total variance (Fig. 2F–I). Factor 1 (42.6% variance) loaded positively on freezing and head scanning and negatively on sniffing, reflecting a defensive–exploratory behavioral axis. Factor 2 (27.1% variance) loaded strongly on rearing and free-air whisking, reflecting active exploratory behavior.

Group comparisons revealed a significant effect of group on Factor 1 scores (F₂,₂₈ = 36.78, p < 0.000001, partial η² = 0.72). Obs-S rats showed significantly higher Factor 1 scores than both Obs-R and Obs-CTL rats (p < 0.001), whereas Obs-R rats did not differ from Obs-Ctrl rats (p = 0.26) (Fig. 2H). Factor 2 scores did not differ between groups (Fig. 2I).

### Mesoscale Organization of cFos Correlation Networks

To gain insights into the organization of functional brain networks associated with fear expression, we employed cFos immunohistochemistry to identify neurons active during OFL (*20*). Based on the fraction of cFos-expressing neurons per brain region, we constructed a fully connected graph in which each brain region analyzed resembles a node and the Pearson’s correlation factor between the cFos ratios of a pair of brain regions determines the weight of the corresponding edge. The resulting Pearson correlation matrices of regional cFos expression across animals within each group were subjected to Voronoi clustering to characterize the mesoscale organization of the underlying brain network. (Fig. 3). This analysis showed clear differences in cluster structure across groups, with control rats showing multiple medium-sized clusters, Obs-S rats dominated by a single large cluster, and Obs-R rats displaying one large cluster together with numerous singleton nodes.

**Figure 3:**
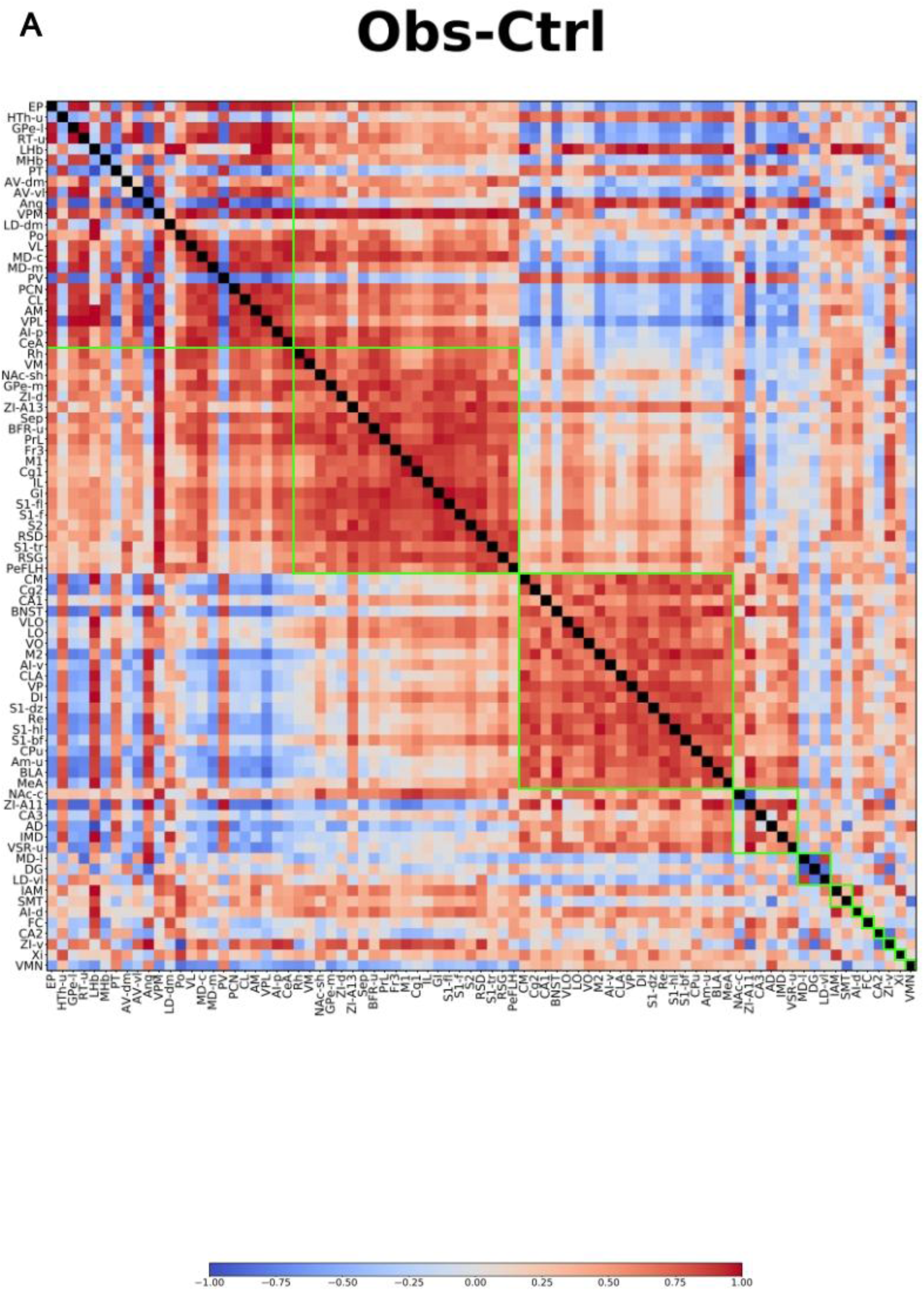
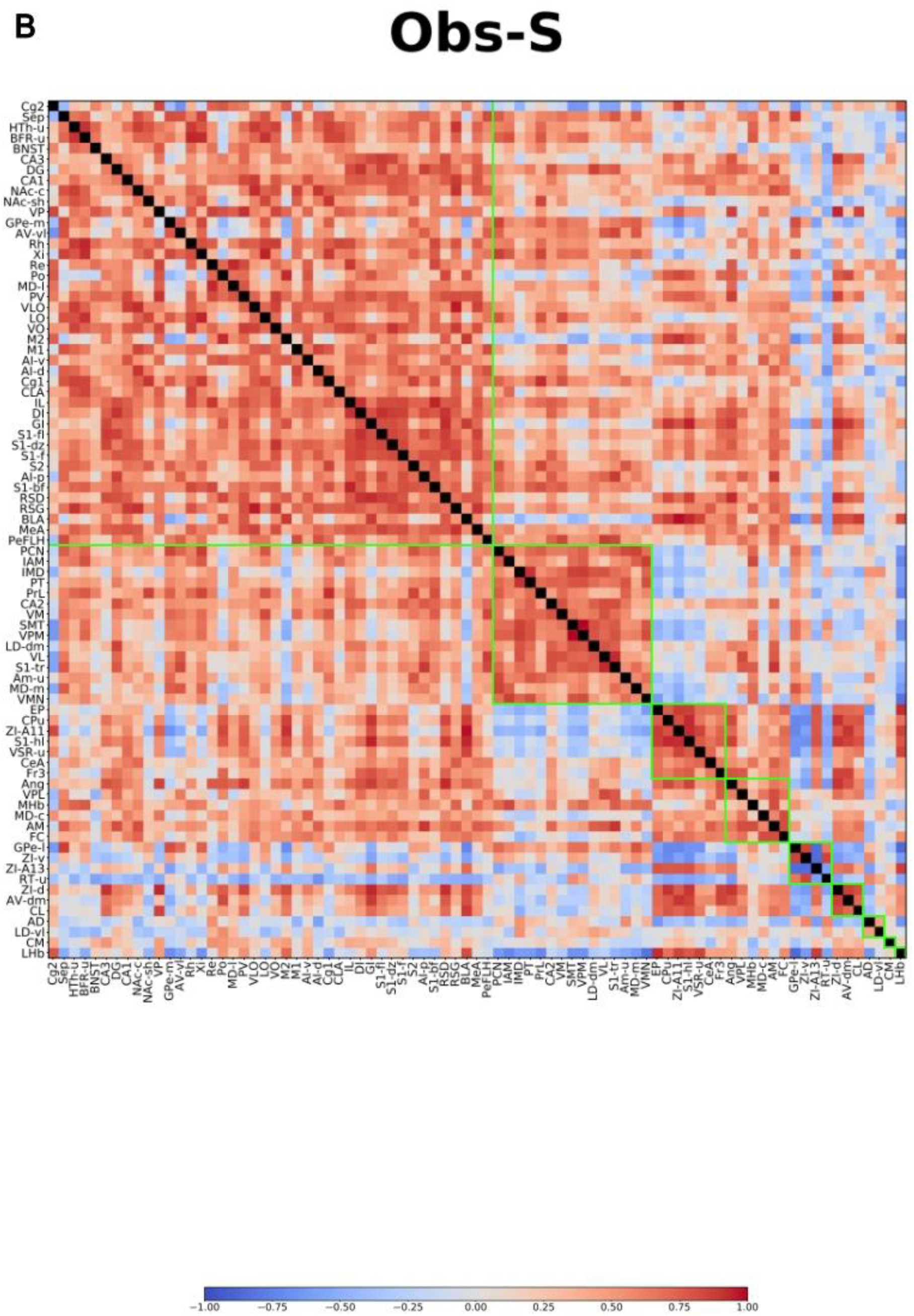
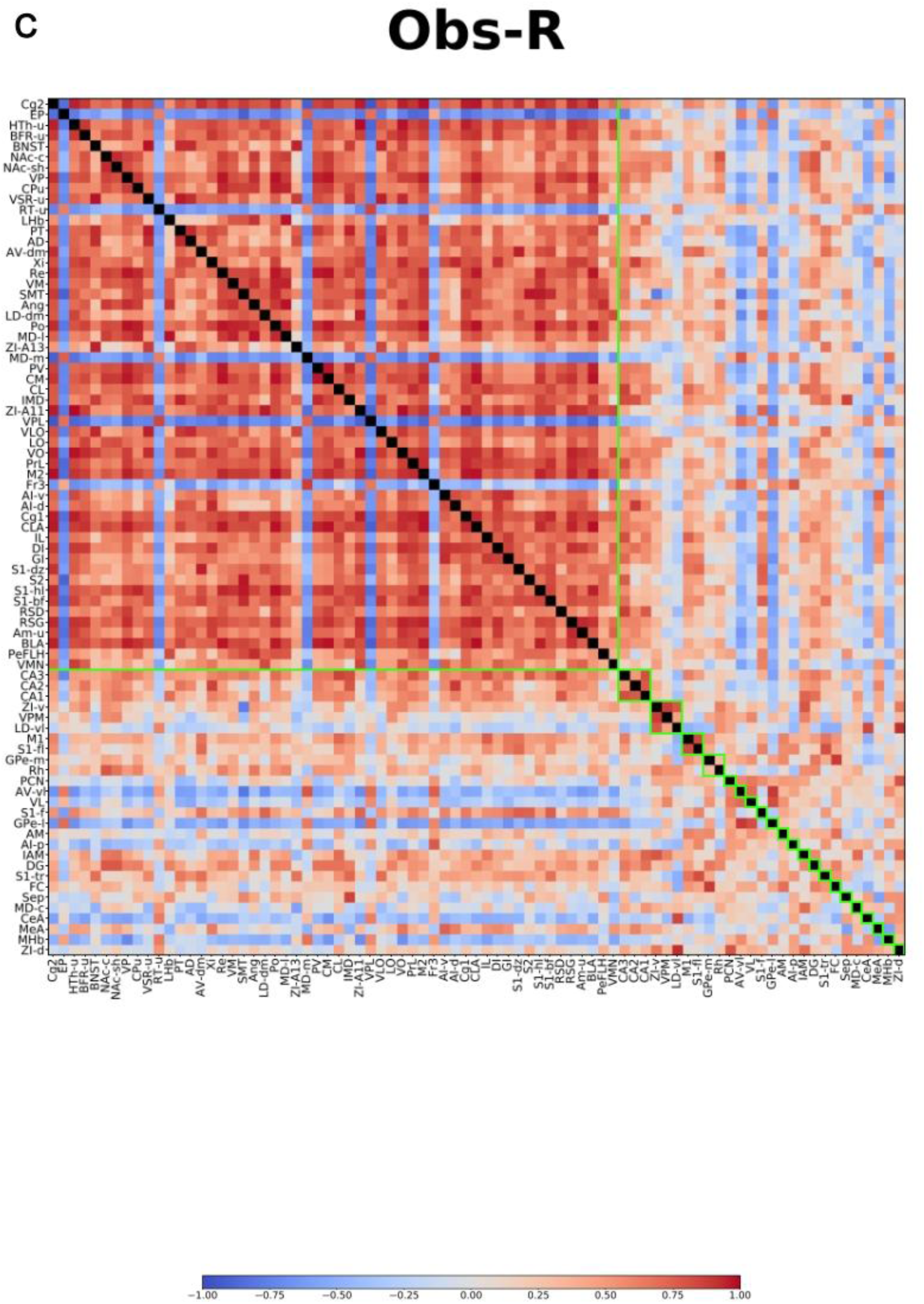
Correlation matrices of c-Fos activity across brain regions following observational fear learning. Correlation matrices showing pairwise Pearson correlations of c-Fos expression across 81 brain regions in control observers (Obs-Ctrl), observational-susceptible rats (Obs-S), and observational-resilient rats (Obs-R). Each matrix represents correlations calculated across animals within each group. Red colors indicate positive correlations and blue colors indicate negative correlations. Colored boxes (green) highlight clusters of correlated regions identified by the Voronoi clustering algorithm. The diagonal represents self-correlations.

In Obs-Ctrl rats, the network partitioned into 12 clusters (6 major clusters and 6 clusters containing a single brain region), including clusters enriched in thalamic and cortical regions. Within these clusters, both positive and negative correlations were present. In Obs-S rats, cluster partitioning was reduced (9: 7 major, 2 single region clusters), with most regions grouped into a single large cluster and only a small number of additional clusters. Correlations within clusters were predominantly positive, but less strong than in OBS-CTL. In Obs-R rats, the network contained one large cluster spanning cortical, thalamic, limbic, striatal, and hypothalamic regions together with a substantially larger number of singleton clusters relative to the other groups. Within the large cluster, both positive and negative correlations were observed. and a structure of subclusters appeared. Moreover, the absolute correlation values within this main clusters were high.

Together, these results show that the three groups differ in both the number of clusters and the distribution of regions across clusters.

### Centrality Profiles Across Groups

To determine whether specific brain regions occupied prominent positions within these networks, node centralities were quantified using betweenness, closeness, and weighted degree centralities. Across centrality measures and group comparisons, several midline and cortical regions—including the claustrum (CLA), mediodorsal thalamus (MD-m), central medial thalamus (CM), and cingulate cortex (Cg2)—showed higher centrality in Obs-R rats relative to Obs-S rats (Fig.6). In Obs-Ctrl rats, several regions exhibited higher centrality relative to Obs-S rats, including the zona incerta (ZI-A13) and secondary somatosensory cortex (S2) based on betweenness centrality (Fig.4). Additional differences between Obs-Ctrl and Obs-R rats involved higher betweenness centrality in granular insular cortex (GI), ZI-A13, and central amygdala (CeA), as well as higher weighted degree in several thalamic regions including the mediodorsal thalamus (MD-c) and anteromedial thalamus (AM) (Fig. 5). No differences in closeness centrality were detected between Obs-Ctrl and Obs-S rats (Fig. 4). Comparisons involving Obs-R rats revealed a distinct pattern of central nodes. Relative to Obs-S rats, Obs-R rats showed higher betweenness centrality in cingulate cortex (Cg2), mediodorsal thalamus (MD-m), central medial thalamus (CM), and CLA (Fig. 6). Closeness centrality was also higher in MD-m. Weighted degree analyses similarly identified increased connectivity of several regions in Obs-R rats, including CLA, Cg2, M2, and thalamic nuclei (Fig. 6). Direct comparison of Obs-R and Obs-Ctrl rats further indicated higher weighted degree in Obs-R networks within cortical and subcortical regions including CLA, Cg2, and ventral orbital cortex (VO) (Fig. 5).

**Figure 4:**
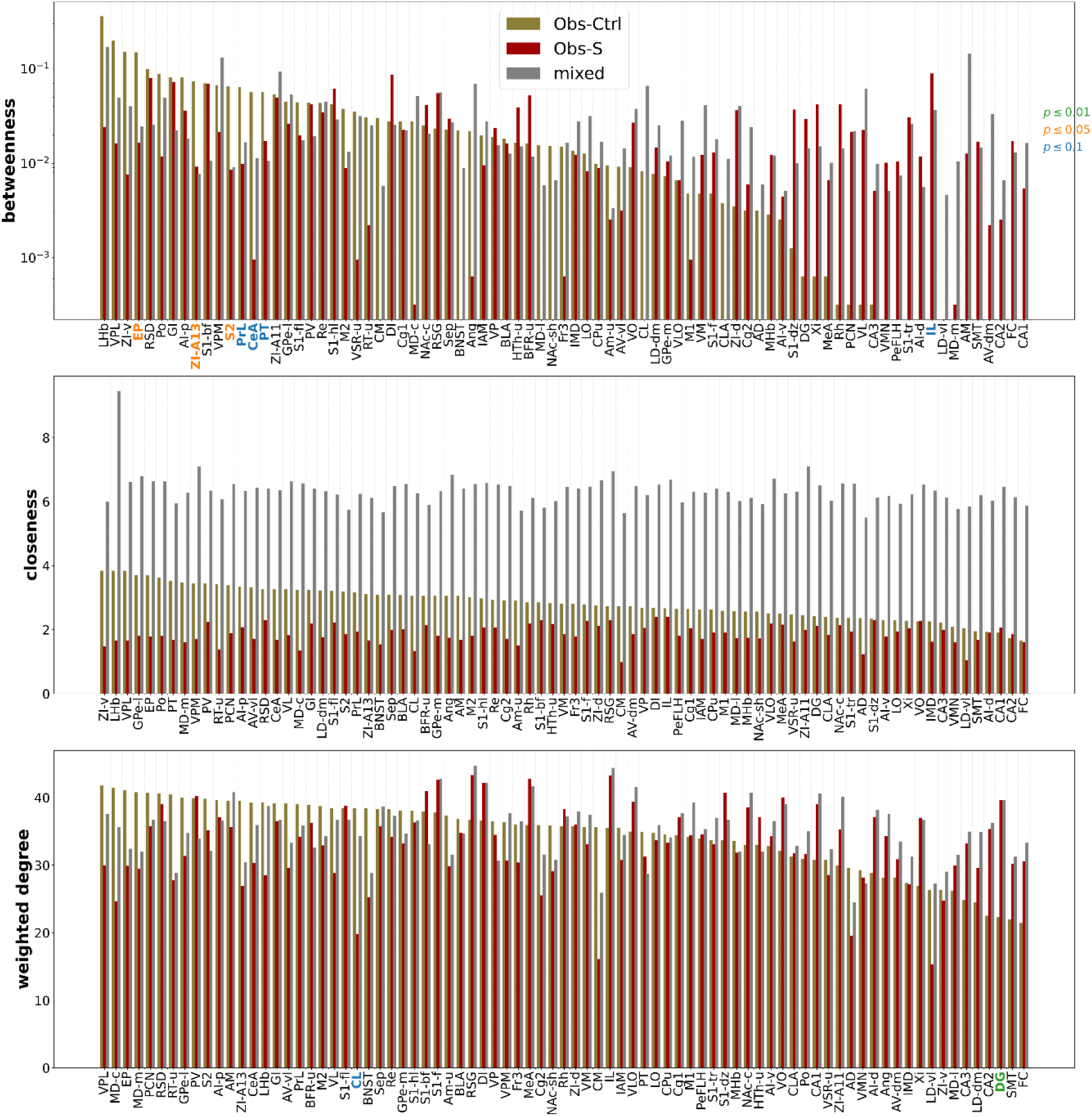
Node centrality measures across brain regions in observational fear networks. Bar plots showing betweenness centrality (top), closeness centrality (middle), and weighted degree (bottom) for each brain region in control observers (Obs-Ctrl), observational-susceptible rats (Obs-S), and the mixed reference network. Regions are ordered along the x-axis according to their centrality values. Colored labels indicate regions showing significant differences between groups based on permutation testing (green, p ≤ 0.01; orange, p ≤ 0.05; blue, p ≤ 0.1).

**Figure 5:**
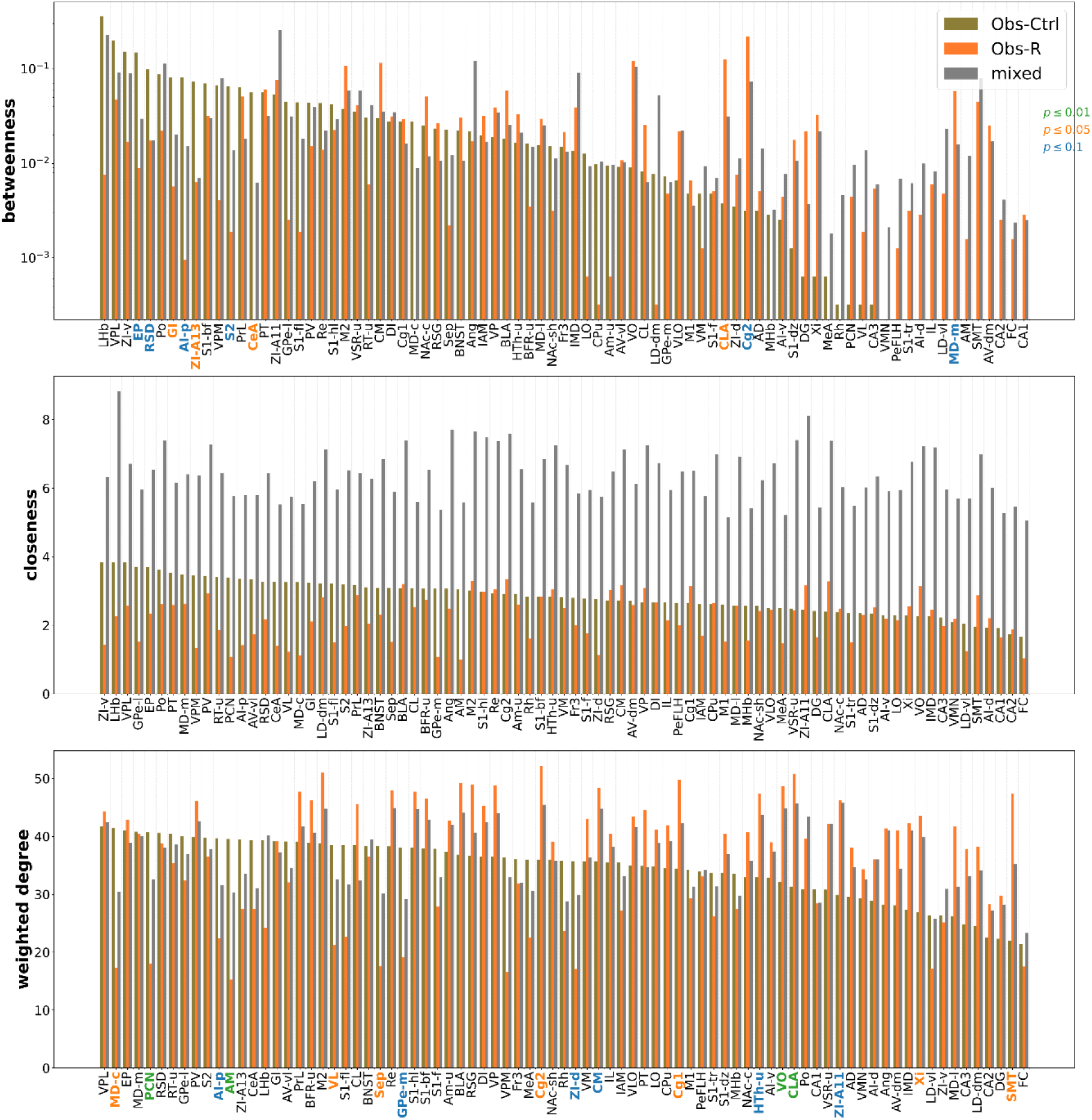
Node centrality measures in control and observational-resilient brain networks. Bar plots showing betweenness centrality (top), closeness centrality (middle), and weighted degree (bottom) for each brain region in control observers (Obs-Ctrl), observational-resilient rats (Obs-R), and the mixed reference network. Regions are ordered along the x-axis according to their centrality values. Colored labels indicate regions showing significant differences between groups based on permutation testing (green, p ≤ 0.01; orange, p ≤ 0.05; blue, p ≤ 0.1).

**Figure 6:**
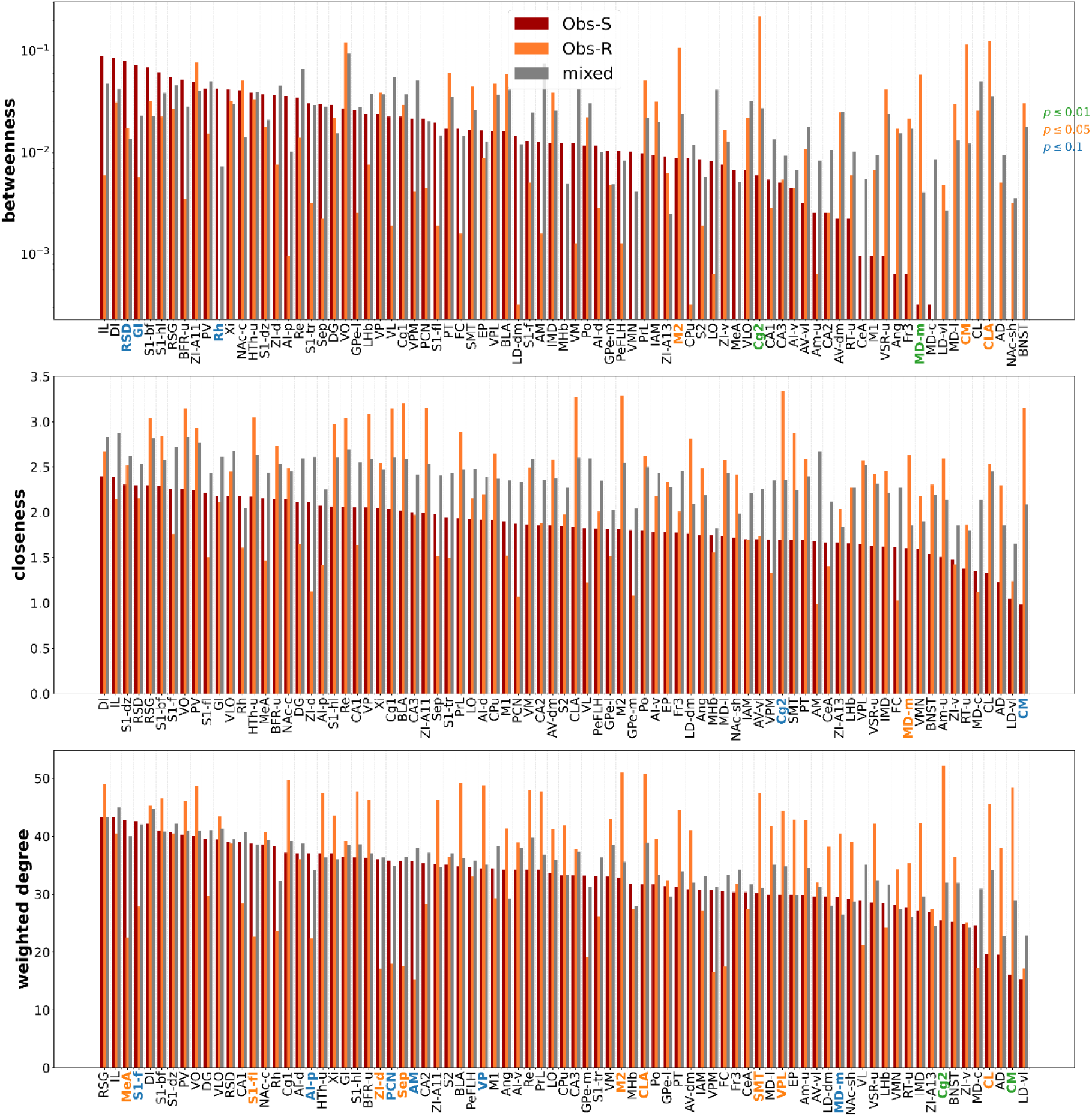
Node centrality measures in observational-susceptible and observational-resilient brain networks. Bar plots showing betweenness centrality (top), closeness centrality (middle), and weighted degree (bottom) for each brain region in observational-susceptible (Obs-S), observational-resilient (Obs-R), and the mixed reference network. Regions are ordered along the x-axis according to their centrality values. Colored labels indicate regions showing significant differences between groups based on permutation testing (green, p ≤ 0.01; orange, p ≤ 0.05; blue, p ≤ 0.1).

Across centrality measures and group comparisons, several regions, including the CLA, MD-m, central medial thalamus CM, and Cg2 consistently emerged as prominent nodes within the functional networks, particularly in Obs-R rats.

### Communication Efficiency Under Progressive Link Removal

To examine how activity might propagate through these networks, global and local communication efficiency were measured during progressive edge removal. This analysis showed group differences primarily during weak-link removal, whereas efficiency trajectories were largely similar when strong correlations were removed first.

Global communication efficiency (Eg) was evaluated by progressively deleting either the strongest or weakest correlations and comparing the resulting efficiency curves with a permutation-based null model generated by random mixing of animals between groups (Fig. 7). When strong correlations were removed first, efficiency trajectories for both, Eg and El, were largely similar across Obs-CTL, Obs-S, and Obs-R rats. In contrast, when weak correlations were removed first, differences between groups became apparent. In the Obs-Ctrl versus Obs-S comparison, global efficiency was slightly lower in Obs-S rats across a restricted range of edge densities. In the Obs-Ctrl versus Obs-R comparison, Obs-R rats showed a larger reduction in Eg across a broader range of densities. Similarly, in the Obs-S versus Obs-R comparison, Eg was lower in Obs-R rats across early and intermediate densities. Local communication efficiency (El) showed the opposite pattern during weak-link removal (Fig. 7). Under weak-link removal, local efficiency was slightly lower in Obs-S rats than in Obs-Ctrl rats across a restricted range of densities, albeit not reaching significance. In contrast, Obs-R rats showed higher local efficiency than both groups across intermediate densities.

**Figure 7:**
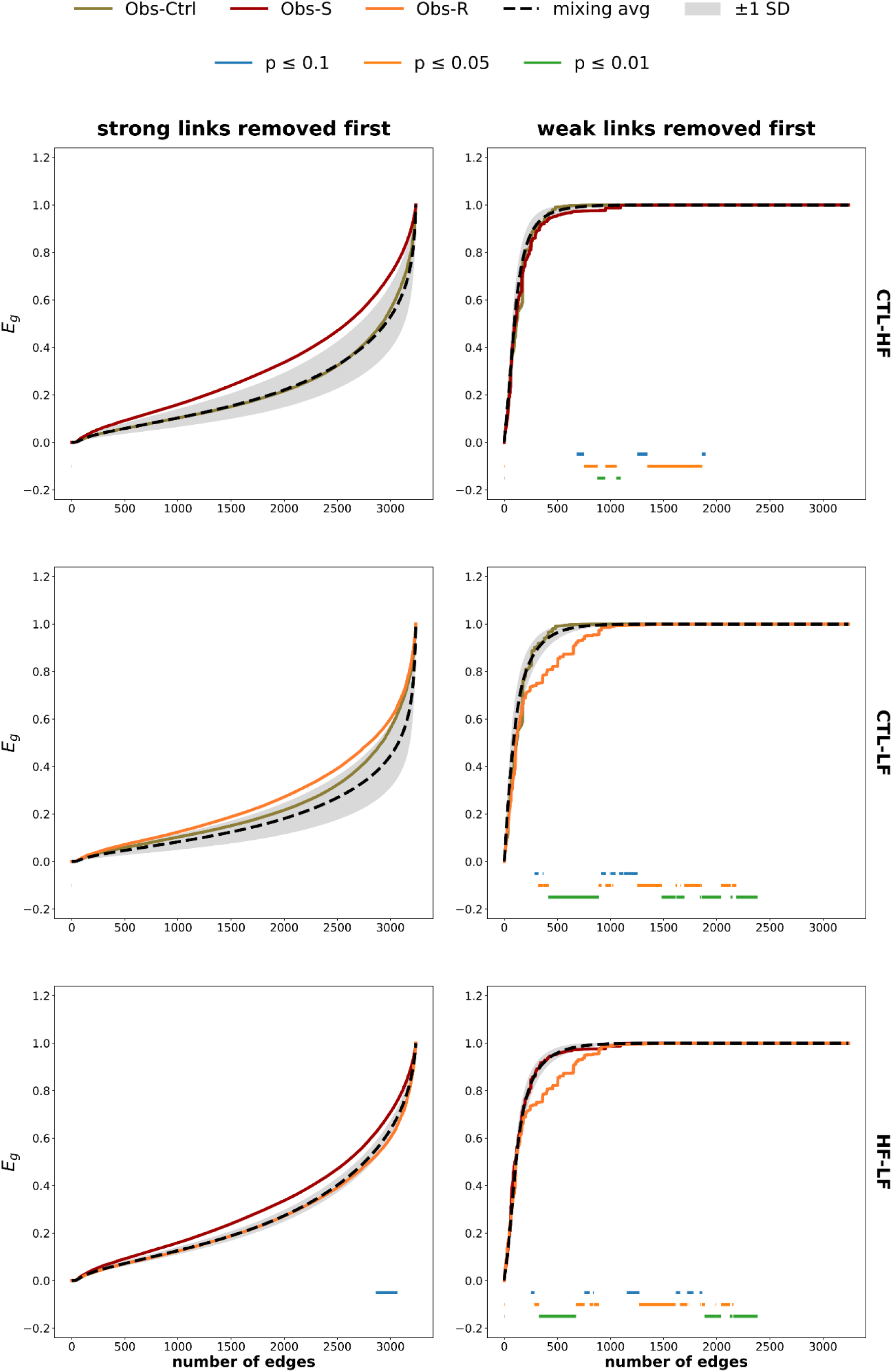
Global communication efficiency during progressive edge removal. Global efficiency of functional networks as edges are progressively removed. Networks from control observers (Obs-Ctrl), observational-susceptible rats (Obs-S), and observational-resilient rats (Obs-R) are compared with a mixed reference network (dashed line; shaded area indicates ±1 SD). The left column shows efficiency curves when strong correlations are removed first, whereas the right column shows efficiency when weak correlations are removed first. Colored horizontal bars indicate ranges of edge densities showing significant differences between networks based on permutation testing (green, p ≤ 0.01; orange, p ≤ 0.05; blue, p ≤ 0.1).

### Backbone Structure and Network Geometry

To visualize the structure of the network and the correlations shaping the network, network backbones were extracted and mapped using forced atlas layouts (Fig. 8,9,10). Backbone representations showed differences in the polarity and spatial distribution of strong correlations across groups, with Obs-S networks dominated by positive correlations but relatively weak and a compact layout (Fig. 9A,B), whereas Obs-R networks retained both strong positive and strong negative correlations with a more spatially dispersed organization (Fig. 10 A,B). In Obs-Ctrl rats (Fig. 8 A,B), the backbone contained both positive and negative correlations linking cortical, thalamic, hippocampal, and striatal regions that form distinct clusters. In Obs-R rats, both positive and negative correlations were present, and the network layout appeared more spatially dispersed. In the Obs-R network (Fig. 10 A,B), the claustrum and mediodorsal thalamus were positioned near the central portion of the layout and connected to multiple edges within the network core.

**Figure 8:**
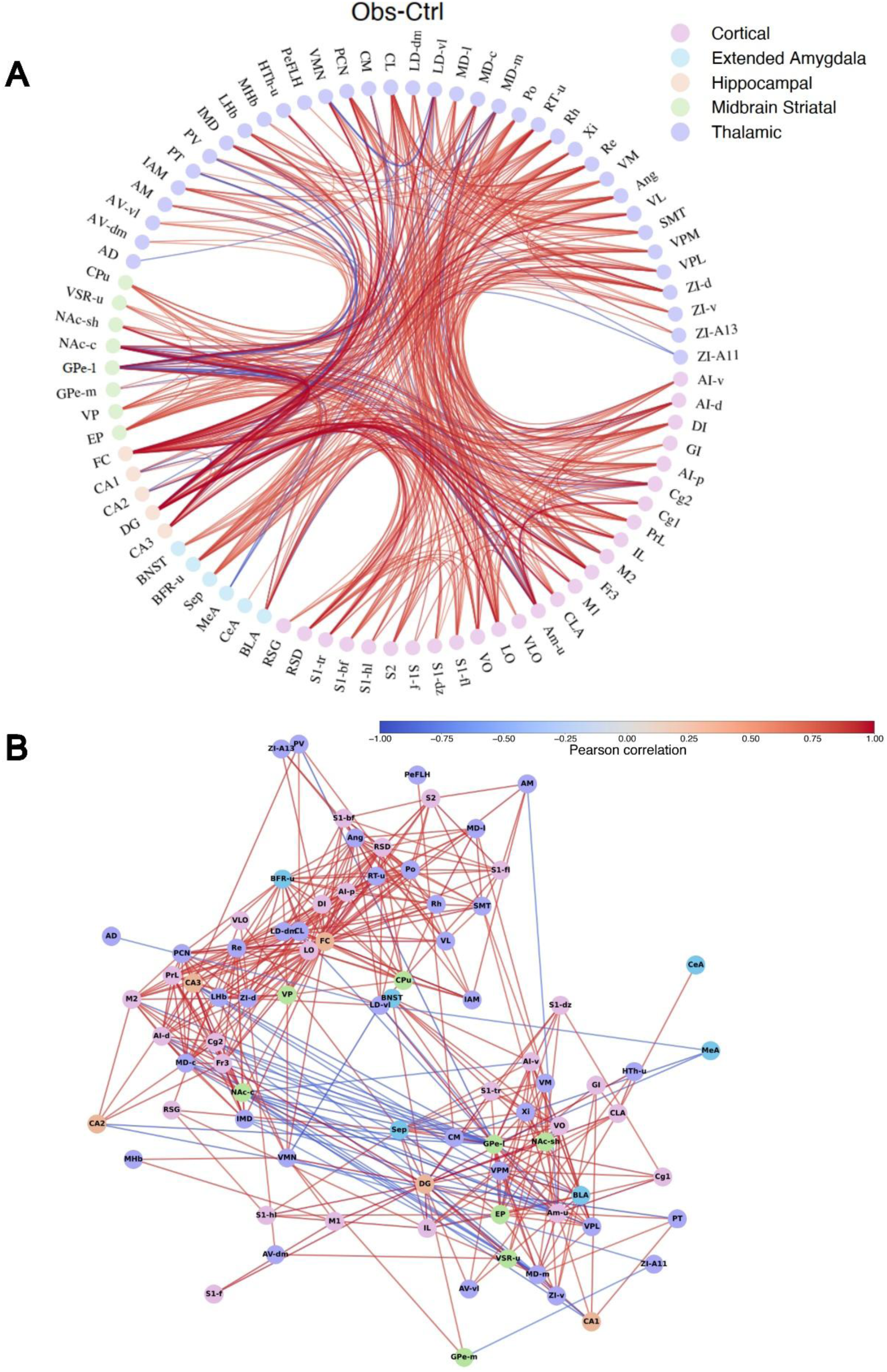
Local communication efficiency during progressive edge removal. Local efficiency of functional networks as edges are progressively removed. Networks from control observers (Obs-Ctrl), observational-susceptible rats (Obs-S), and observational-resilient rats (Obs-R) are compared with a mixed reference network (dashed line; shaded area indicates ±1 SD). The left column shows efficiency curves when strong correlations are removed first, whereas the right column shows efficiency when weak correlations are removed first. Colored horizontal bars indicate ranges of edge densities showing significant differences between networks based on permutation testing (green, p ≤ 0.01; orange, p ≤ 0.05; blue, p ≤ 0.1).

**Figure 9:**
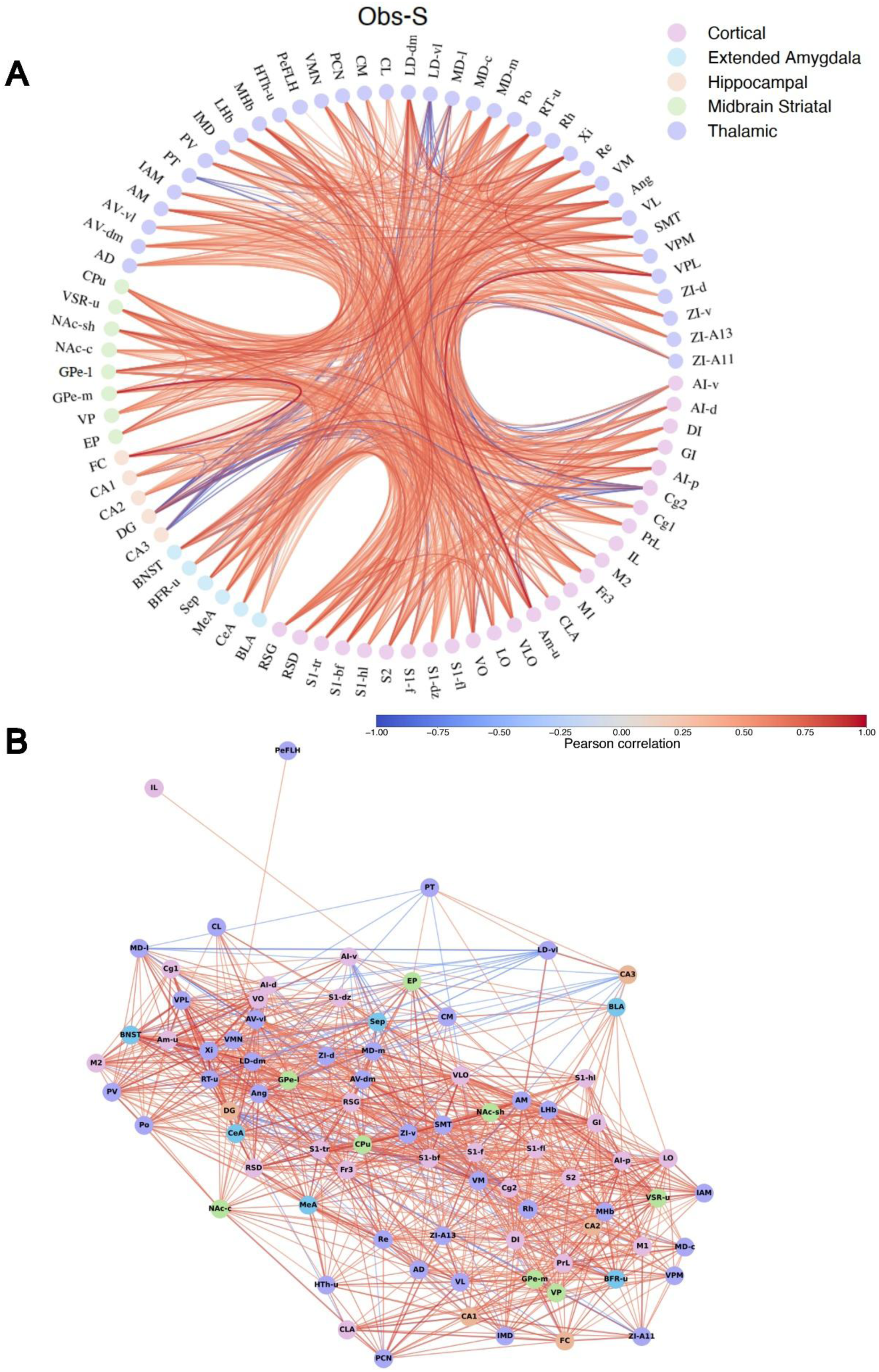
Network backbone representation of c-Fos correlations in control observers. (A) Circular representation of the functional network derived from Pearson correlations of c-Fos expression across brain regions in control observers (Obs-Ctrl). Nodes represent brain regions and are grouped by anatomical system (color-coded). Edges represent correlations between regions, with red indicating positive correlations and blue indicating negative correlations. (B) Force-directed layout of the network showing the backbone structure of the strongest correlations. Nodes correspond to brain regions and edges represent significant correlations between regions. Node colors indicate anatomical groupings as in panel A.

**Figure 10:**
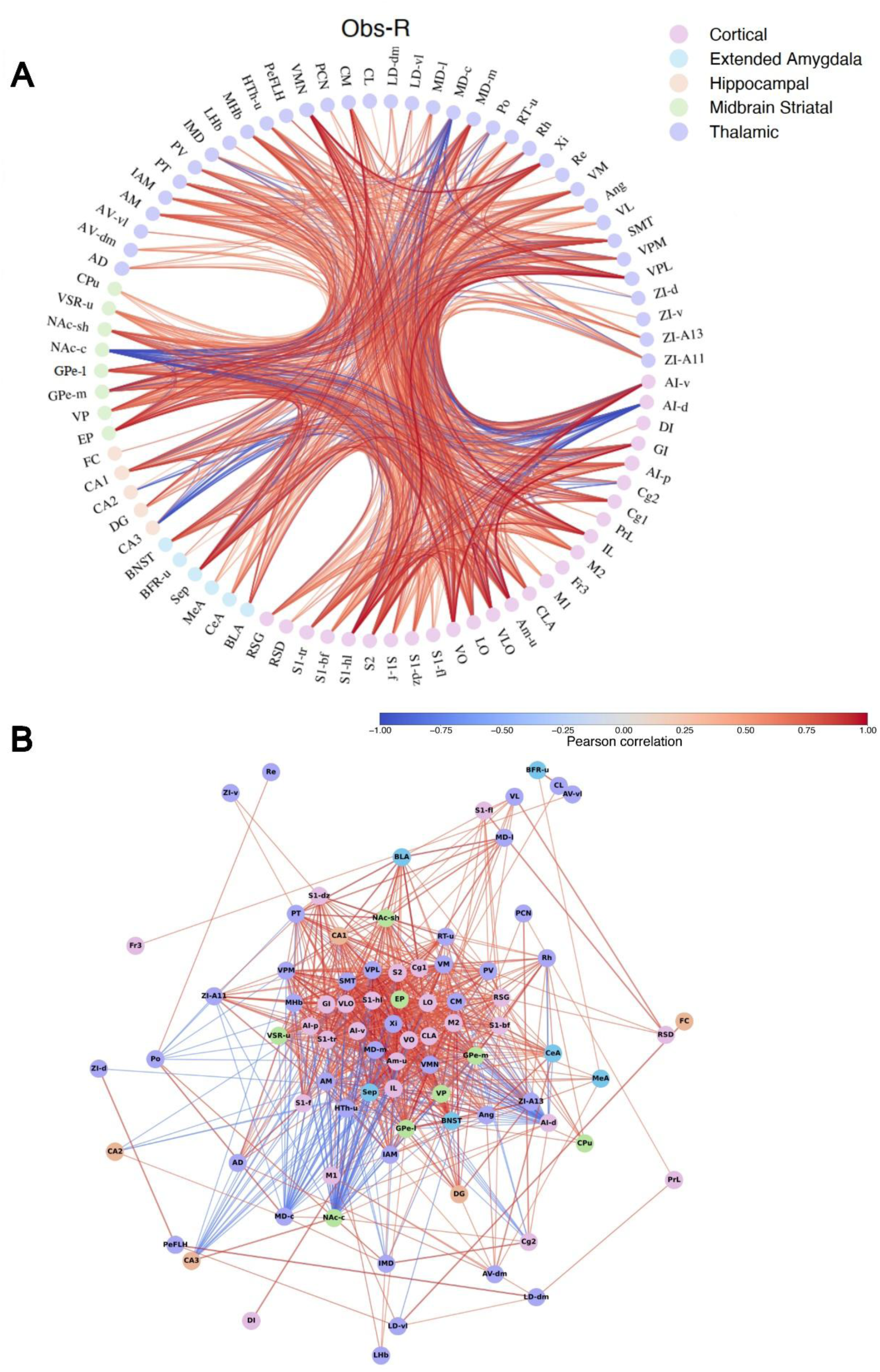
Network backbone representation of c-Fos correlations in observational-susceptible rats. (A) Circular representation of the functional network derived from Pearson correlations of c-Fos expression across brain regions in observational-susceptible rats (Obs-S). Nodes represent brain regions and are grouped by anatomical system (color-coded). Edges represent correlations between regions, with red indicating positive correlations and blue indicating negative correlations. (B) Force-directed layout showing the backbone structure of the strongest correlations in the Obs-S network. Nodes correspond to brain regions and edges represent significant correlations between regions. Node colors indicate anatomical groupings as in panel A.

## DISCUSSION

In the present study, we examined how socially acquired fear is represented across distributed brain networks and whether individual differences in behavioral responses to OFL are associated with differences in network organization. By combining behavioral phenotyping with brain-wide cFos mapping across 81 regions and graph-theoretical analysis, we found that OFL does not produce a uniform response across observer rats. Instead, animals segregated into distinct behavioral phenotypes that were associated with different configurations of large-scale brain networks. These findings suggest that variability in socially acquired fear is associated with differences in the organization of distributed neural networks, rather than with the engagement of entirely distinct brain regions.

Behaviorally, OFL exposure produced two subpopulations: a majority of observer rats that exhibited no freezing or freezing levels that did not exceed those observed in control animals (Obs-R) and a smaller subset that expressed robust freezing (Obs-S). Ethological analysis further showed that these differences were captured by a dominant defensive-exploratory behavioral axis, along which Obs-R rats clustered with control rats. Importantly, this behavioral divergence occurred despite evidence that both subpopulations detected the OFL experience as biologically significant. Although only Obs-S rats displayed overt defensive behavior, both Obs-S and Obs-R rats showed elevated corticosterone during OFL acquisition and expression. This indicates that both groups registered the socially transmitted threat and formed the underlying association. This pattern argues against a simple interpretation in which Obs-R rats failed to detect or encode the socially conveyed fear. Instead, the results indicate that individual differences reflect variation in the regulation of socially transmitted fear information, influencing whether it is ultimately expressed as defensive behavior. One factor that may contribute to this behavioral distribution is the relatively mild nature of the OFL procedure used here. Our protocol included only six cue-shock pairings, whereas recent mouse OFL studies have used substantially larger numbers, including approximately thirty pairings in the study by Silverstein and colleagues (*9*). Within a threat-imminence framework (*21*), lower perceived threat levels are expected to keep animals in a pre-encounter or low-imminence state, characterized by minimal defensive responding and preserved exploration, whereas higher perceived threat levels are required to elicit overt post-encounter behaviors such as freezing. This framework provides a plausible explanation for why most observers in the present study did not exhibit freezing beyond control levels, despite endocrine evidence that the OFL cue was biologically salient.

Consistent with this behavioral heterogeneity, large-scale network analyses showed phenotype-specific differences in the organization of correlated activity across brain regions. Control observers exhibited a modular network structure composed of several medium-sized clusters containing both positive and negative correlations, consistent with previously reported patterns of modular organization in functional brain networks derived from c-Fos activity mapping (*22*). In contrast, networks of susceptible animals were characterized by reduced cluster partitioning and increased global synchronization, with most regions grouped into a single large cluster dominated by positive correlations. Resilient animals displayed a different configuration consisting of one large cluster together with numerous singleton nodes. Together, these patterns suggest that socially acquired fear is associated with differences in how brain regions become functionally grouped into coordinated modules, with susceptible animals showing stronger global coupling and resilient animals maintaining a more differentiated network structure. This pattern aligns with prior work showing that c-Fos–derived functional networks during fear memory retrieval exhibit greater segregation than degree-matched random networks, indicating that modular organization reflects structured network states rather than a statistical artifact (*23*).

Differences between phenotypes were also reflected in the topological roles of specific regions within the networks. Several regions, including the claustrum, cingulate cortex, and midline thalamic nuclei such as the mediodorsal and central medial thalamus, occupied more central positions within the networks of resilient animals. These structures are anatomically well positioned to link cortical, limbic, and subcortical systems(*24–26*), suggesting that resilient observers may rely on network configurations in which integrative nodes play a more prominent coordinating role. In contrast, networks of susceptible animals showed higher centrality primarily in cortical and hippocampal regions. These results indicate that differences in behavioral responses to socially acquired fear are associated with shifts in the influence of particular nodes within distributed networks rather than the recruitment of entirely distinct circuits.

Network efficiency analyses further indicated that the phenotypes differed in how correlations contributed to large-scale communication across the system. Differences between groups emerged primarily when weaker correlations were removed, suggesting distinct relationships between global integration and local clustering. Under these conditions, global communication efficiency decreased from controls to susceptible animals and further to resilient animals, whereas local efficiency showed the opposite pattern. This opposing relationship suggests that weaker correlations contribute differently to large-scale integration and local clustering across the networks, potentially reflecting differences in how distributed neural systems coordinate responses to socially conveyed fear.

Visualization of network backbones provided a complementary perspective on these organizational differences. Networks of susceptible animals were dominated by positive correlations and appeared more compact, whereas resilient networks retained both positive and negative correlations and exhibited a more spatially distributed structure. Within the resilient network, the claustrum and mediodorsal thalamus occupied central positions in the backbone and connected to multiple edges within the network core, consistent with their elevated centrality values. Together, these observations suggest that these regions occupy influential positions within the networks of observer rats displaying minimal defensive behavior.

These findings extend prior work demonstrating that fear memories are embedded within distributed brain networks rather than localized circuits. In particular, studies combining cFos mapping with graph-theoretical analysis have shown that contextual fear memory engages brain-wide networks containing influential hub regions whose disruption can impair memory consolidation (Vetere et al., 2017). The present results extend this framework to socially acquired fear by showing that differences in behavioral responses are associated with differences in the organization of distributed networks supporting emotional processing. Rather than reflecting simple differences in activation of canonical fear structures, variability in OFL responses appears to emerge from differences in how these regions are embedded within broader network architectures.

Several limitations should be considered. Network organization was assessed at a single post-learning time point, and future studies will be required to determine how these configurations evolve over time or relate to longer-term behavioral outcomes. In addition, while cFos mapping provides a brain-wide snapshot of activity patterns across multiple regions, it does not capture temporal dynamics or causal interactions between nodes. All experiments were conducted in male rats, and future work will be needed to determine whether similar network phenotypes are observed in females, given well-documented sex differences in stress responses and defensive behavior. Combining large-scale activity mapping with causal perturbation approaches targeting candidate hub regions will be important for determining whether the network nodes identified here play functional roles in shaping individual responses to socially acquired fear.

In summary, observational fear learning produces heterogeneous behavioral responses among observer rats that are accompanied by distinct large-scale network architectures. By linking behavioral phenotypes to differences in distributed network organization, the present findings provide a systems-level framework for understanding how susceptibility and resilience to socially acquired fear may emerge from variability in the connectivity of large-scale neural systems.

## Ethics statement

Procedures were approved by the Local Animal Ethics Committee at Linköping University and were in accordance with the EU Directive 2010/63/EU on the protection of animals used for scientific purposes as implemented in Swedish national regulations.

## Author contributions

Conceptualization: NM, KC, EB; Data curation: NM, BM, JR, KC, AL, LX, LM; Formal analysis: NM, BM, JR, KC, AL; LM; Funding acquisition: MER; CK, EB Investigation; NM, KC, AL, EB; Methodology: NM, BM, JR, KC, AL, MER; CK, EB; Project administration: NM, BM, JR, MER; CK, EB; Software: NM, BM, JR; Supervision: MER; CK, EB; Validation: MER; CK, EB; Visualization: NM, BM, JR; Writing original draft: NM, BM, JR, EB; Writing – review & editing: NM, BM, JR, MER; CK, EB.

All authors contributed to the article and approved the submitted version.

## Acknowledgments

The authors thank the Center for Biomedical Resources (CBR) at Linköping University for the functioning of all laboratory animal facility and the caring of laboratory animals. Figures were created with BioRender.com.

